# GTA-5: A Unified Graph Transformer Framework for Ligands and Protein Binding Sites — Part I: Constructing the PDB Pocket and Ligand Space

**DOI:** 10.64898/2026.02.27.708634

**Authors:** Bogdan C. Ciambur, Raphaël Pageau, Olivier Sperandio

## Abstract

Structural recognition between a protein target and a ligand underpins therapeutic innovation, yet computational representations of protein binding sites and small molecules remain largely disjoint. Here we introduce GTA-5, a unified graph transformer auto-encoder framework designed to capture the geometric structure and chemical composition of ligands and protein binding pockets, embedding them into multidimensional latent spaces where proximity reflects functional compatibility. Ligands and pockets are represented as three-dimensional point clouds annotated with *Tripos* atom type labels, omitting explicit bond connectivity to enable structural reasoning based on spatial context rather than predefined connectivity graphs. By not enforcing bond topology, GTA-5 maintains representational flexibility across molecular modalities while preserving chemically meaningful local environments. The model was trained on a curated dataset from the Protein Data Bank comprising 64,124 liganded pockets and 23,133 unique ligands spanning 2,257 protein families. We find that functional protein families cluster coherently in both pocket and ligand latent spaces while retaining biologically meaningful heterogeneity. The model captures physicochemical pocket properties such as volume, exposure, and hydrophobicity directly from raw structural data, while ligands with distinct scaffolds co-localise when occupying similar binding environments. This provides a basis for several downstream applications including scaffold hopping in ligand-based virtual screening, QSAR/QSPR modelling using embedding-derived descriptors, and drug repurposing via pocket similarity. More broadly, the GTA-5 framework establishes a foundation for structural reasoning across molecular modalities in drug discovery.

## 1 Introduction

Drug discovery is ultimately a problem of navigating structure. Therapeutic innovation depends on understanding how molecular shape, chemical composition, and spatial organization govern recognition between ligands and biological targets. Recent advances in structural biology and artificial intelligence, including high-accuracy protein structure prediction methods such as *AlphaFold* (Jumper et al., 2021), have dramatically expanded access to three-dimensional molecular data. Yet computational representations of this structural universe remain fundamentally fragmented. Small molecules are typically encoded as molecular graphs using message-passing neural networks (MPNNs, e.g., Gilmer et al. 2019, Liu et al. 2023) or transformer-based architectures such as *GROV ER* (Rong et al., 2020) and *Graphormer* (Ying et al., 2021). In contrast, protein binding sites are described through voxel-based convolutional neural networks as in early 3D CNN scoring models, or through handcrafted cavity descriptors implemented in tools such as *V olSite* Desaphy et al. (2012), *ProBiS* Konc and Janežič (2010), or *FINDSITE* Brylinski and Skolnick (2008). As a consequence, models that perform well within a single representational paradigm rarely generalize across structural modalities, with notable exceptions being *AlphaFold*3 (Jumper et al., 2023), *Boltz*-2 (Huang et al., 2023), and *RFDiffusion* (Trippe et al., 2023), which nonetheless focus on the accurate modeling of specific complexes (rather than broad-scale datasets) and do not natively handle small molecules or ligands within the same (protein-focused) architectural framework.

This fragmentation limits our ability to reason about transferability. Scaffold definitions formalized by Bemis and Murcko (1996) and scaffold-hopping strategies described by Brown and Jacoby (2006) highlight the importance of identifying structural compatibility beyond conventional chemical similarity. Structure-based drug repurposing approaches such as *FINDSITE* demonstrate that binding-site similarity can transcend evolutionary relationships. However, when ligands and binding pockets are embedded in incompatible representational spaces, structural comparison becomes indirect and heuristic. A unified structural language capable of encoding diverse molecular entities within a common geometric and semantic space remains largely absent.

Here we introduce GTA-5, a Graph Transformer Autoencoder framework designed to address this representational divide. Rather than encoding molecules and pockets as fundamentally different graph objects, GTA-5 adopts a modality-agnostic abstraction: molecular objects are represented as labeled three-dimensional point clouds. This design is inspired by geometric deep learning approaches for unordered spatial data, such as *PointNet* (Qi et al., 2019), and extends ideas from three-dimensional equivariant architectures including *SchNet* (Schütt et al., 2018) and *SE*(3) − *Transformers* (Fuchs et al., 2022). Each point is defined solely by spatial coordinates and a categorical chemical label. Explicit bond connectivity, central to conventional molecular graph neural networks, is intentionally omitted, though partly accounted for implicitly through the choice of chemical labels. This choice shifts the representational emphasis from fixed chemical topology to spatially contextualized chemical identity, enabling ligands and binding cavities to be processed under a single architectural paradigm. While canonical chemical graphs encode how atoms are bonded, molecular recognition is ultimately governed by three-dimensional spatial organization. Models such as *SchNet* and *SE*(3) − *Transformers* have demonstrated the importance of explicitly modeling geometry in molecular learning tasks. Using proximity-based graphs built dynamically from spatial neighborhoods, rather than predefined bonds, GTA-5 captures local chemical environments and long-range structural dependencies without constraining the model to topology-specific assumptions.

The model is trained in a self-supervised manner through geometric and semantic reconstruction, following principles introduced in latent molecular representation learning frameworks such as the variational autoencoder approach of Gómez-Bombarelli et al. (2018). The resulting latent embeddings position each object—ligand or binding pocket—within continuous structural manifolds where proximity reflects shared geometric and compositional compatibility rather than predefined (e.g., molecular fingerprint) similarity. This work represents a step toward unified latent spaces in which ligands, pockets, and potentially peptides, can be embedded within a shared geometric language, enabling structural transferability in drug discovery.

In this study, we focus on respectively encoding protein binding pockets and their small-molecule ligands as a foundation for this broader objective. We demonstrate that a modality-agnostic, attention-based geometric framework can recover interpretable structural organization directly from raw three-dimensional data. The remainder of the article is structured as follows. In Section 2 we describe the compilation, curation, and annotation of the core training datasets, including pocket detection and ligand processing. Section 3 details the GTA-5 model architecture, data representation, and training. In Section 4 we present the resulting pocket and ligand latent spaces, highlighting emergent functional clustering. Finally, Section 5 discusses the implications, limitations, and future directions of this approach, situating GTA-5 as a generalizable framework for unified structural representation in drug discovery.

## 2 Data

### 2.1 Retrieval and Quality Control

The dataset used to train our model was compiled following the protocol described in Moine-Franel et al. (2024), hereafter referred to as M24. Briefly, the entire Protein Data Bank (Berman, 2000) metadata (as of April 2025) was retrieved from the Protein Data Bank in Europe (PDBe)^1^ repository, and the annotations within were used to identify protein-ligand complexes as follows. Complexes were required to contain at least one small molecule with a minimum of five heavy atoms, all necessarily drug-like (i.e., C, N, O, S, P, F, Cl, Br, I, and B). This primary list was first pruned by discarding ligands such as ATP, co-factors, or molecules originating from crystallization buffers, due to their limited relevance towards drug design. A series of quality filters were next applied to only retain 3D structures determined by nuclear magnetic resonance, X-ray crystallography (imposing that the resolution ≤ 3.5 Å and |*R*_*free*_ − *R*_*factor*_| ≤ 0.07) or cryogenic electron microscopy (cryo-EM, imposing that the resolution ≤ 3 Å and Fourier shell correlation ≤ 0.143). The remaining structures were downloaded and processed further – incomplete amino acids in the structures were repaired using *FoldX* (Schymkowitz et al. 2005, version 5) and all of the protein-ligand complexes were protonated with the OPLS-AA force field of *GROMACS* (Spoel et al. 2005, version 2020), and saved in *Tripos mol*2 format.

### 2.2 Pocket Detection

To detect and characterise the liganded binding pockets in our dataset we employed the *IChem* suite’s *V olSite* algorithm (Desaphy et al., 2012). For each complex the ligand was used as a reference for the selection of surrounding residues defining the binding site cavity. *V olSite* “fills” this cavity with pseudo-atoms, or probes, placed on a regular 3D grid with 1 Å spacing. Each pseudo-atom is assigned a *Tripos* atom type depending on its local chemical environment given by its nearest protein residues. The options available for these *Tripos* labels are shown in Table 1. It is worth stating that, for *V olSite* pockets, these atom types do not correspond to physical atoms as they would in a molecule, but are only an arbitrary mapping of various properties (e.g., aromaticity, hydrophobicity, etc.) resulting in a 3D pocket object consistent with the *Tripos mol*2 file format, which is nevertheless particularly convenient for our own purposes.

**Table 1:**
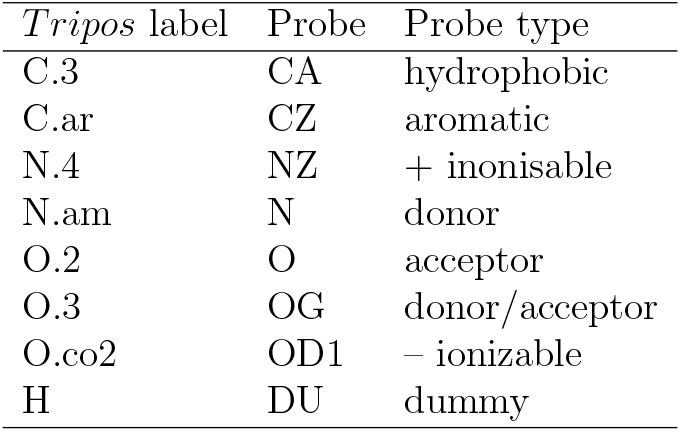
The *Tripos* atom types that may be assigned to *V olSite* pocket pseudo-atoms, to characterise local chemical environment.

*V olSite* was run on all structures with adjusted parameters following M24, producing a set of diverse 3D pocket objects rich in chemical but also geometric information. Global pocket descriptors, such as hydrophobicity, exposure, volume, asphericity, etc., may be analytically calculated from these objects and further used to define an *N*-dimensional ‘pocket-space” (with *N* being the number of descriptors), as was done by M24 (see also Mareuil et al. 2024) for the *pocketome* of protein-protein interactions. We have calculated the same set of 109 *V olSite* and *RDKit* pocket descriptors as M24 to serve as annotations and support model interpretability, though (deliberately) not for any model training per se. Indeed, the goal of this work is for GTA-5 to *learn* the crucial features of these protein-ligand complexes agnostically, in an unsupervised and general manner, *directly* from the raw data, i.e., 3D pocket objects and ligands in their bio-active conformation. Where available, structures were further annotated with their respective entries from the Pfam protein families database^2^ (Mistry et al., 2021).

In total the final dataset comprised 45,666 structures spanning 2,257 Pfam domains (with 1,422 structures lacking a Pfam annotation), with a total of 64,124 liganded pockets, and 23,133 unique ligands. Figure 1 shows the volume distribution of pockets, while the most prevalent protein families are displayed in Figure 2. The obtained pocket and ligand files stored in *mol*2 format served as the training data for GTA-5 − pockets and ligands are treated as 3D point clouds, as we describe in detail in the following Section.

**Figure 1.**
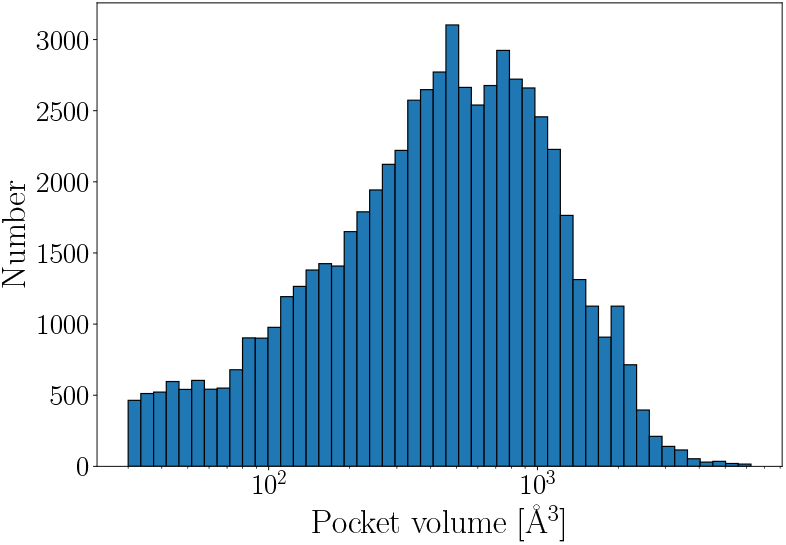
The volume distribution of *V olSite*-detected pockets.

**Figure 2.**
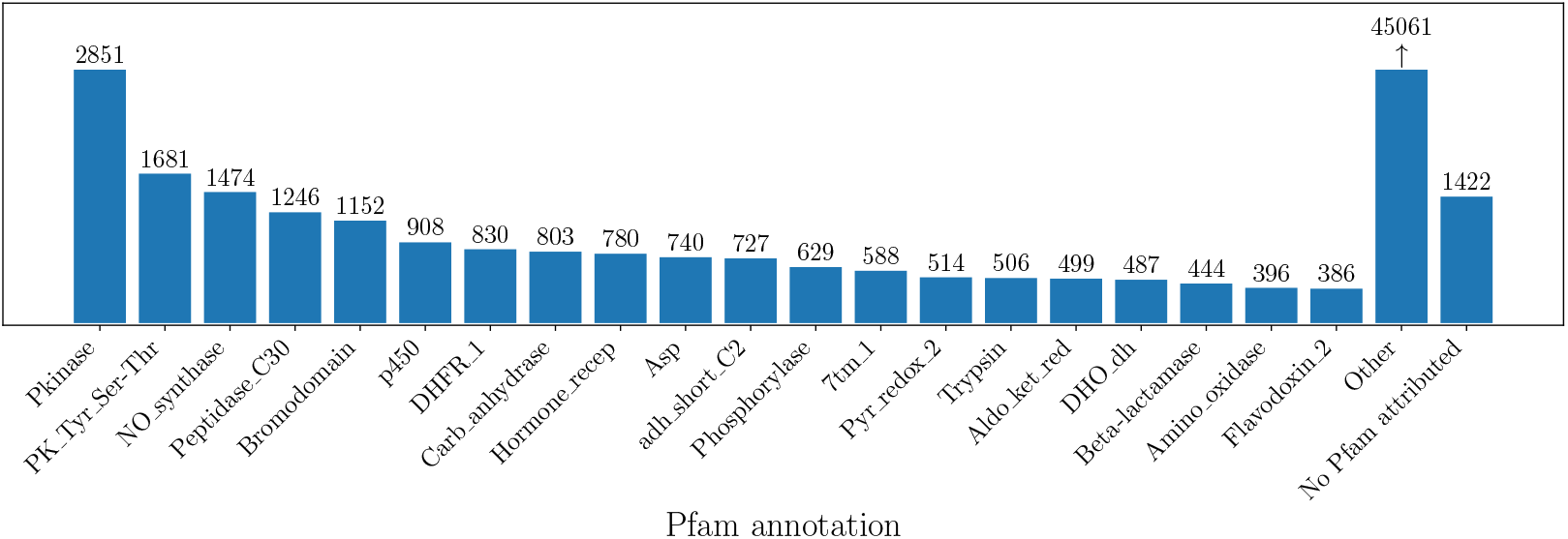
The 20 most commonly-occurring Pfam domains in our final dataset.

**Figure 3.**
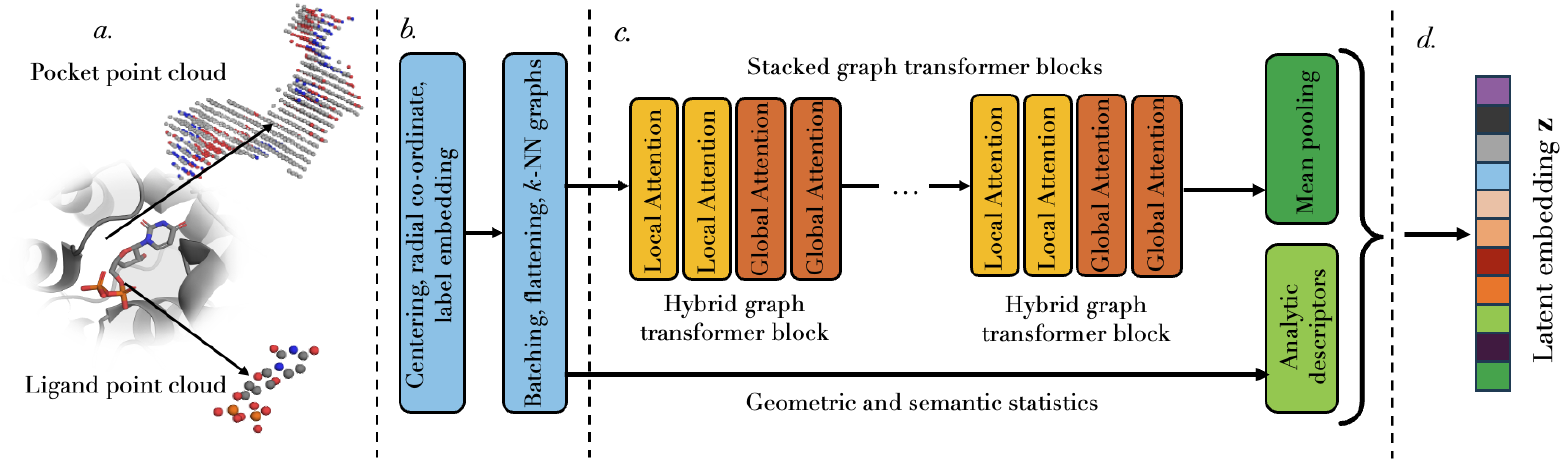
Schematic view of the GTA-5 encoder architecture. The input data (panel *a*.), either small molecules or pocket objects, is represented as 3D point clouds labelled by *Tripos* atom types. The point clouds are pre-processed (panel *b*.) and passed to stacked graph transformer blocks, generating point-wise embeddings aggregated per cloud by mean pooling. In parallel, independent geometric and semantic statistics are concatenated to the pooled feature. This produces the final latent embedding of the point cloud (panel *d*.), which is the primary output of the encoder.

## 3 The GTA-5 Model

GTA-5 is an unsupervised Deep Learning method designed to learn directly from different molecule objects, be it protein binding sites or small molecule binders (or indeed peptide binders, as we shall explore in future work), and encode them into compact and semantically rich latent vectors that capture their 3D geometric structure and semantic composition (i.e., their chemistry, as given by *Tripos* atom type annotation). The model was explicitly designed to be robust to the different types of input data considered in this study, treating all objects as 3D point clouds, where each point (atom or pseudo-atom, accordingly) is characterised solely by its (*xyz*) co-ordinates and *mol*2 atom type label. Atomic bond information, where applicable, is deliberately omitted in order to retain model flexibility and ensure its operability under a single common framework.

### 3.1 Model Architecture

GTA-5 is a hybrid graph transformer auto-encoder (GTA) which leverages local (sparse) and global (dense) attention mechanisms to account for both immediate neighbourhood structure and semantics, as well as long-range interactions within point clouds. This learned relational representation additionally integrates an explicit, analytically calculated set of geometric descriptors (akin to M24, e.g. volume, principal axes, anisotropy, etc.) to finally produce a latent representation as a fixed-sized vector. During training, the model is optimized to reconstruct the original point cloud from this latent representation via an encoder-decoder algorithm. In downstream use, only the trained encoder and the embedding vector are used, the decoder’s role being solely to provide an unsupervised training objective.

#### 3.1.1 Input Data Representation

The fundamental input data instance is a cloud 𝒫 of *N* points represented as

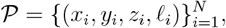

where (*x*_*i*_, *y*_*i*_, *z*_*i*_) ∈ ℝ^3^ are spatial coordinates of point *i*, and *ℓ*_*i*_ ∈ {0, …, *C* − 1} is a discrete label obtained from categorically embedding the set of *Tripos* atom types that occur in the training set, which constitute the model’s label vocabulary. As shown in Table 1 above, *C* = 8 for *V olSite* pockets, whereas for the small molecules in the training set *C* = 24. As the number of points *N* varies across samples, clouds are padded to a common size per batch and accompanied by a binary mask to indicate which entries correspond to valid points.

For such tasks where only intrinsic geometry is informative, absolute position in space constitutes nuisance variation and may hinder generalization. As such, the point clouds are centered, ensuring translation invariance and a stable computation of global statistics. For cloud *j*:

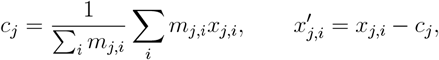

where *m*_*j,i*_ is the validity mask which excludes padded elements. After centering, the radial distance of each point to the geometric centre is computed as

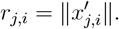

This scalar quantity is rotation-invariant, and combined with the subsequent use of Euclidean distance for neighborhood construction, this provides a representation that does not depend on global orientation.

Finally, semantic information is incorporated through a learned embedding of categorical labels. Each label *ℓ*_*j,i*_ is mapped via a trainable embedding function

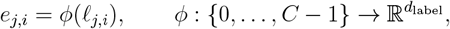

where *ϕ* is implemented as a learnable lookup table. Rather than one-hot embedding the labels, this representation allows the embedding space geometry to adapt during training, enabling classes that frequently co-occur, or play similar geometric roles, to acquire similar representations.

The concatenation of the radial feature and label embedding is projected into a hidden feature space of dimension *d*_hidden_, and forms the initial node representation 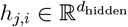.

#### 3.1.2 Sparse Attention – Local Reasoning

To model local spatial relationships, a graph is constructed independently within each point cloud. For each point *i*, its neighborhood 𝒩_*k*_(*j, i*) is defined as the set of indices corresponding to the *k* nearest neighbors in Euclidean distance. In practice, graph construction is performed in a batch-aware manner. All valid points across a batch are temporarily concatenated into

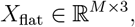

where *M* is the number of non-padded points in the batch as given by the validity mask. A batch-constrained k-nearest neighbor operation is then applied to *X*_flat_, ensuring (through a companion vector storing the parent cloud index of each point) that neighbors are selected only among points belonging to the same cloud.

Given initial node features *h*_*j,i*_, local attention updates each point by aggregating information exclusively from its neighborhood. For each cloud *j*, learned linear projections produce query, key, and value representations:

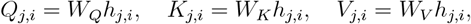

where *W*_*Q*_, *W*_*K*_, *W*_*V*_ are the usual trainable parameters. From these, attention weights are computed in the usual form from the surrounding neighbours *m* ∈ 𝒩_*k*_(*j, i*) of each node, i.e.:

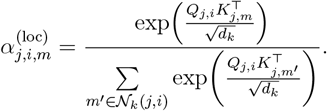

The locally updated feature is then

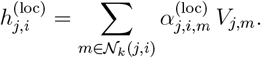

This sparse attention mechanism allows each node to refine its representation by gaining awareness of its closest neighbours. Information flows along the kNN graphs akin to message-passing neural networks, and indeed the higher the depth of the model, the further out a given node’s awareness reaches. However, unlike many graph-based molecular representations that employ attention-based message passing, here we do not impose a fixed graph connectivity (e.g., the atomic bonds of a molecule) but rather make use of a purely proximity-based graph representation. This ensures that the model is general enough to train on both molecule and pocket objects, respectively, and allows a certain degree of flexibility in its applications, as we explain further in the Discussions section below.

#### 3.1.3 Global Reasoning – Dense Attention

While local neighbourhoods capture fragmentary structure in a given molecular object, it is necessary to also capture overall shape and chemical composition through long-range interactions, particularly when dealing with binding pockets. To enable such global reasoning, dense self-attention is computed, enabling every node to become aware of every other node within each cloud.

For cloud *j*, attention is evaluated across all valid point pairs (*i, i*^*′*^) with 1 ≤ *i, i*^*′*^ ≤ *N*_*j*_. Using the same query, key, and value projections as above, global attention weights are defined as

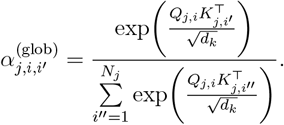

The globally aggregated representation is

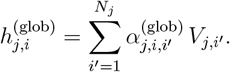

This dense attention mechanism allows distant but structurally or semantically related points to influence each other, enabling the model to capture global shape dependencies that cannot be inferred from purely local neighborhoods.

#### 3.1.4 Graph Transformer Blocks

The outputs of local and global attention are concatenated and projected:

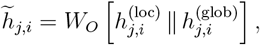

where 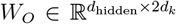 is a learned projection. Each hybrid transformer block operates on the collection of node features *H* = {*h*_*j,i*_}. Given normalized inputs LayerNorm(*H*), the hybrid attention module computes 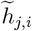 for each point and applies point-wise the first residual update,

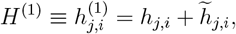

A position-wise two-layer feedforward network with GELU activation and dropout, is then applied independently to each node,

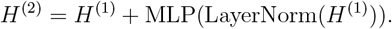

Since attention is computed within clouds and the MLP acts independently per node, permutation equivariance within each cloud is preserved. The model stacks a depth of *L* = 4 such blocks to gain progressively refined point-wise representations.

#### 3.1.5 Explicit Global Descriptors

The representations emerging from the graph transformer blocks are complemented by a set of global, analytically calculated and independent descriptors to enrich the final latent representation. For each cloud *j*, global geometric descriptors are computed from the centered coordinate tensor *X* ∈ ℝ^*N ×*3^. The covariance matrix

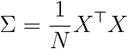

yields eigenvalues *λ*_1_, *λ*_2_, *λ*_3_, describing the principal axes of variance. Together with axis-wise variances, bounding-box volume, the eigenvalues of the inertia matrix *I*_1_, *I*_3_, *I*_3_, and the index

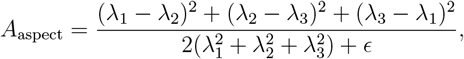

which measures eigenvalue disparity and so shape anisotropy, these form an 11-dimensional geometric descriptor *g*. The descriptor is normalized using a running mean and variance across batches, producing

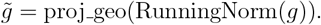

Additionally, semantic composition is captured by class frequencies *p*_*c*_ and label entropy *H*:

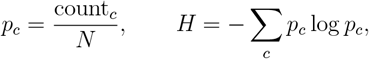

which are concatenated into a vector *ℓ* and normalized via running statistics:

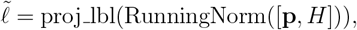

yielding a compact semantic embedding. The running normalization is updated each epoch and ensures stability across batches as the training progresses.

#### 3.1.6 Latent Embedding

After the final hybrid transformer block, the resulting point-wise embeddings *h*_*j,i*_ are aggregated per cloud *j* by mean pooling:

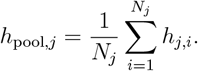

The pooled feature is concatenated with the projected geometric and semantic descriptors described above, and mapped through a linear layer:

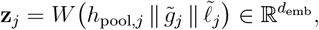

where *d*_emb_ is the desired (tunable) dimensionality of the latent space. This yields the final global embedding of cloud *j* and constitutes the primary output of the encoder.

#### 3.1.7 Decoder and Training Objective

To train the encoder without external supervision, a decoder reconstructs the original point cloud from the latent embedding **z**. The embedding is broadcast to all nodes, projected to the hidden dimension, and refined through another stack of hybrid transformer blocks. Two linear heads predict reconstructed coordinates 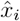 and label logits 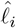.

The loss over valid nodes 𝒱 is

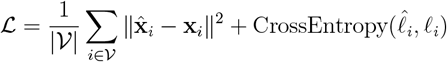

The first term enforces geometric reconstruction fidelity, while the second preserves semantic consistency. This objective encourages the latent embedding to retain information necessary to recover both structure and labels.

### 3.2 Training and Inference

The model is trained in a self-supervised autoencoding regime. Given a batch of point clouds 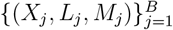, the encoder produces latent representations 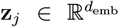, which are passed to the decoder to reconstruct point coordinates and categorical labels at each valid node. Let 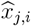 and 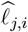 denote the predicted coordinates and label logits, respectively. The loss ℒ is evaluated only over valid points, and model parameters are optimized accordingly using Adam with learning-rate scheduling. During training, the running normalization statistics used for the global geometric descriptor 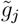 and the semantic descriptor 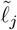 are updated on the fly and stored together with the best-performing model checkpoint, for use in inference mode.

The principal architectural hyperparameters are the embedding dimension *d*_emb_, the hidden feature dimension *d*_hidden_, the number of attention heads *H*, the number of transformer blocks *L*, and the kNN neighborhood size *k*. In addition, the fraction of heads allocated to global attention determines the balance between sparse local and dense global interactions. Optimization hyper-parameters include the learning rate, batch size, maximum number of epochs, learning-rate decay schedule, and early-stopping patience.

At inference time, only the encoder is used. For a new cloud *j*, coordinates and labels are mapped to tensors (*X*_*j*_, *L*_*j*_, *M*_*j*_) using the fixed vocabulary derived from the training set. The stored normalization statistics are restored, and the embedding is computed deterministically as

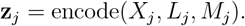

The resulting vector 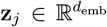 constitutes the final fixed-dimensional representation of cloud *j* and is stored for downstream use.

## 4 Results

### 4.1 Training the GTA-5 Encoder

GTA-5 was trained on the sets of ligands and *V olSite* pockets described in Section 2, with each set producing its own trained encoder. The model was run with a choice of *k* = 16 for kNN graph construction, a hidden feature space dimension of *d*_hidden_ = 256, and a depth of *L* = 4 stacked hybrid transformer blocks each with 4 attention heads split equally between sparse (local) and dense (global) attention. The output embedding dimension was chosen to be *d*_emb_ = 256, sufficiently large to allow for a rich representation yet compact enough to facilitate subsequent distance evaluations and minimum spanning tree construction, as we shall detail further below.

In each case the training was performed with a batch size of 64 over an initially chosen 1000 epochs, however in practice both models converged after ∼ 500 - 600 epochs, after which the patience (100 epochs) was reached, triggering early stopping. As mentioned before, training is unsupervised and optimizes the encoder’s latent representation of these objects, **z**, such that it captures the essential information required to reconstruct the point cloud geometric and semantic structure. As such, the best model checkpoints and running statistics were subsequently used in inference mode to embed these full datasets into respective latent spaces, and explore the way these high-dimensional spaces are structured in order to leverage relative proximity within them as a means to accomplish downstream drug discovery tasks.

### 4.2 The *Pocketome* and *Ligandome*

To visualize the structuring of the pocket and ligand latent spaces as learned by their respective encoders, and assess them as drug discovery tools, we construct a *pocketome* and a *ligandome* similarly to M24, but using our latent embeddings in stead of analytically calculated descriptors as feature vectors.

We first compute pairwise Euclidean distances between all pocket, and respectively ligand, embeddings. The resulting distance matrices may be represented as complete graphs, where nodes represent our objects of interest and edges represent separation (distance) in their respective latent space. To simplify this representation into a more intuitive form of visualization, we generate minimum spanning trees using Kruskal’s algorithm (Kruskal 1956), and display these using the TMAP tool (Probst and Reymond, 2020). The resulting *pocketome* and *ligandome* are shown in Figure 4, both annotated by the corresponding Pfam domain of the parent protein.

**Figure 4.**
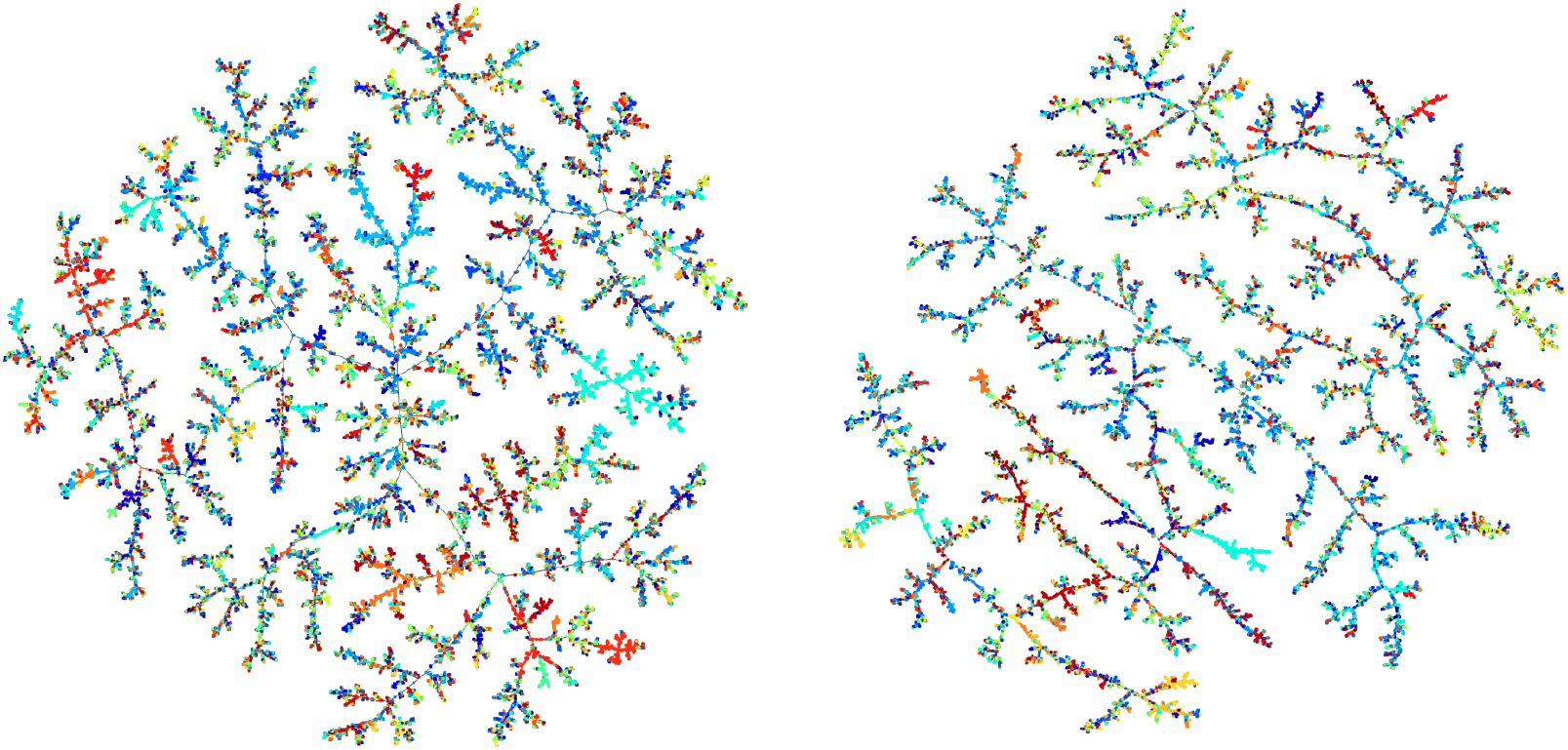
Minimum spanning tree representations of the *pocketome* (left) and *ligandome* (right), constructed from GTA-5 pocket and ligand embeddings, respectively, and colour-coded by Pfam domain.

The pocketome reveals pronounced clustering of pockets from the same Pfam domain, indicating that binding sites are consistently conserved and thus making latent-space proximity a reliable predictor of functional and chemical similarity. This property is particularly relevant for drug repurposing, as pockets that co-localise on the same branch in the GTA-5 pocketome can bind ligands with distinct chemical scaffolds, as we show in Figure 6. As we have seen in Section 2, multiple distinct pockets often occur within a single protein structure. This implies that protein families can subdivide into separate local clusters in this multidimensional pocket space, which we indeed observe to be the case. Interestingly, pockets from different Pfam domains occasionally co-localize in latent space, highlighting opportunities for ligand transferability and drug repurposing, where molecules targeting one pocket could interact with structurally (and functionally) similar pockets in unrelated proteins.

The independent pocket annotations previously calculated with *V olSite* (Section 2.2) confirm that GTA-5 embeddings implicitly capture key physicochemical properties such as volume, hydrophobicity, and solvent exposure. As shown in Figure 5, these features are coherently reflected in the minimum spanning tree, despite the model being trained solely on raw structural data.

**Figure 5.**
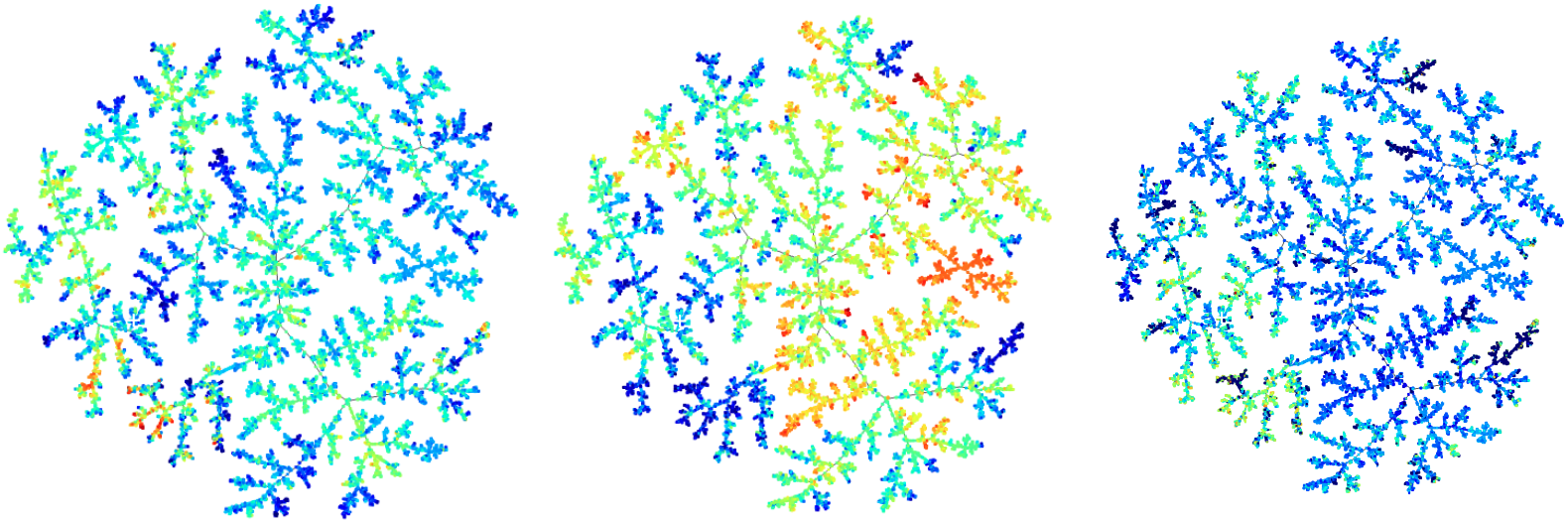
The GTA-5 *pocketome* colour-coded by three pocket annotations calculated independently with *V olSite*, from left to right: hydrophobicity (CA), volume (in logarithmic scale), and pocket exposure (T50-60). The minimum spanning tree is structured coherently with these layers, indicating that GTA-5 is able to capture these pocket properties in its latent representation.

**Figure 6.**
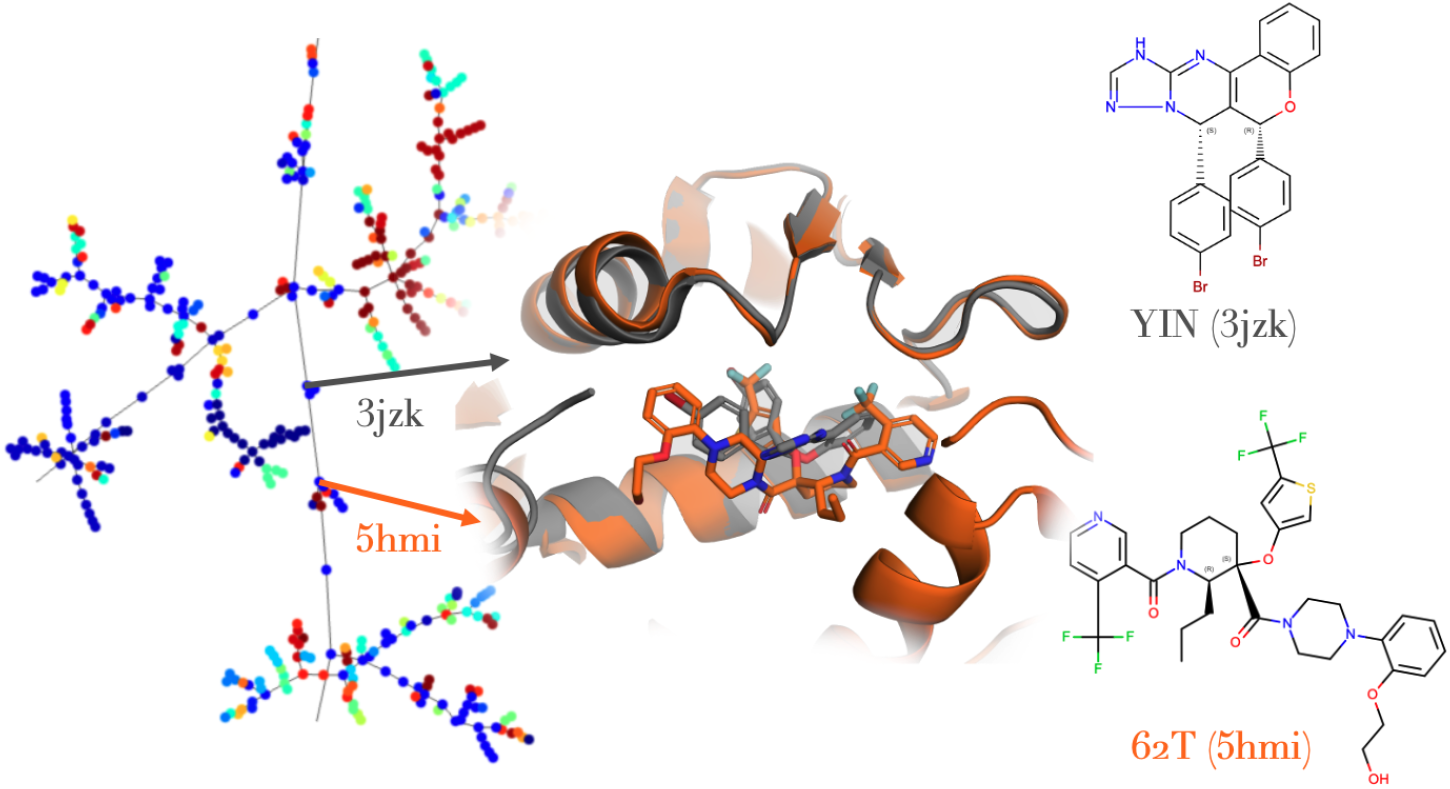
An example of close neighbours in pocket space from the SWIB domain (PDB codes 3jzk and 5hmi), each hosting ligands with very different chemical scaffolds (ligand codes YIN and 62T, respectively).

Surprisingly, the ligandome too displays similar trends. Ligands occupying pockets from the same Pfam domain tend to cluster in latent space, even when their chemical scaffolds differ (Figure 7). Although this clustering is somewhat less pronounced than in the pocketome, this organization facilitates scaffold hopping in ligand-based virtual screening, enabling the identification of novel chemotypes compatible with the binding site through ligand-based virtual screening, taking the bio-active molecule as the reference.

**Figure 7.**
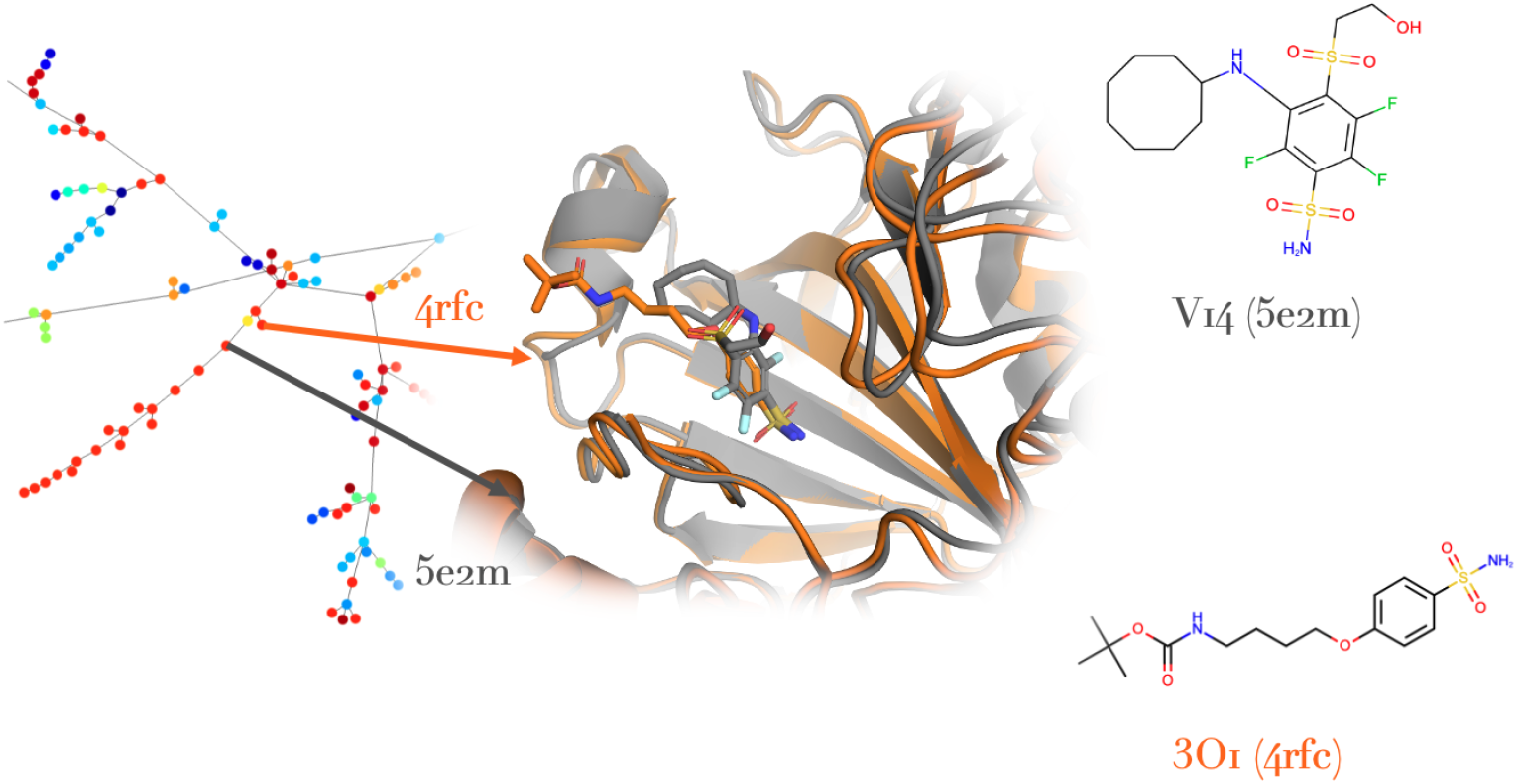
An example of different chemotypes col-localising on the GTA-5 ligandome. Both systems belong to the Eukaryotic-type carbonic anhydrase Protein family, with PDB (ligand) codes 3O1(4rfc), V14(5e2m).

While minimum spanning trees are efficient for visualisation purposes and provide a global and intuitive appreciation of latent space structuration, they nevertheless omit (by construction) significant information and are additionally constrained to connect each point to the common tree, sometimes to the detriment of other otherwise nearer neighbours of the same branch. As such, in the following we assess the clustering of our pocket and ligand spaces with quantitative metrics computed from the full datasets.

### 4.3 Latent Space Structure Metrics

Given embedding vectors **z**_*i*_ with associated Pfam annotation *P*_*i*_, we evaluate how well the latent space is organized by functional family using neighborhood statistics. For each pocket/ligand *i*, we define the kNN purity as the fraction of its *k* nearest neighbors in embedding space, 𝒩_*k*_(*i*), that share its Pfam label:

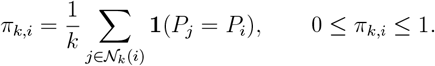

Averaging over all samples *n* (dataset size, see Section 2) yields the global purity,

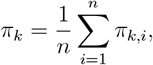

while averaging over proteins belonging to the same Pfam defines a per-class purity.

Because Pfam families are imbalanced in our dataset, as seen in Figure 2, we compare *π*_*k*_ to the random baseline expected under no clustering. If *ν*_*c*_ denotes the global frequency of class *c*, then the expected purity under random mixing is

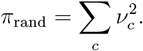

We therefore report a normalized purity

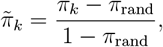

for which 0 corresponds to random mixing and 1 to perfectly pure neighborhoods.

We additionally compute a kNN class entropy statistic. With *p*_*ic*_ denoting the fraction of neighbours of pocket/ligand *i* associated to a Pfam class *c*, the neighborhood entropy is

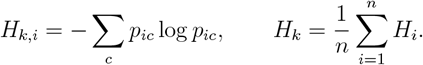

Under random mixing, the expected entropy equals the Shannon entropy of the global class distribution,

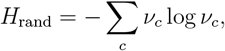

where *ν*_*c*_ again denotes the global frequency of class *c*. We define the normalized entropy reduction,

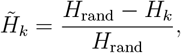

for which 0 indicates random mixing and 1 indicates perfectly pure neighborhoods. Low entropy (or high normalized entropy reduction) reflects well-separated Pfam clusters in latent space.

For our GTA-5 pocket space we obtain a mean Pfam normalised purity of 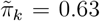 and normalised entropy 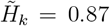, for local neighbourhood size *k* = 10. Together these indicate that the neighbourhoods are concentrated and not always dominated by a single class. This is in part expected biologically but importantly it indicates that the model captures functional similarity, not just strict family boundaries, making it particularly interesting in the context of drug repurposing. We do note that several Pfam classes achieve high purity, particularly well represented domains like Peptidase C30 (0.95, 1246 members), NO synthase (0.93, 1474 members), or the Bromodomain (0.84, 1152 members), which highlights the benefit for the encoder to be exposed to many examples of a certain category.

For our ligandome we report an expectedly lower though still relatively elevated latent space clustering by protein family domain, namely a mean normalised purity value of 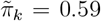 and normalised entropy 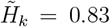, again for local neighbourhood size *k* = 10. As before, high purity values were achieved for more and less well represented Pfam domains, namely TP methylase (0.95, 155 members), Bacteriorhodopsin-like proteins (0.92, 337 members), or NO synthase (0.91, 1474 members).

## 5 Discussion

The central contribution of GTA-5 is not solely architectural but representational. By encoding protein binding pockets and ligands within a common three-dimensional point-cloud formalism, we demonstrate that structurally distinct molecular entities can be embedded into coherent latent landscapes without reliance on predefined connectivity graphs or handcrafted similarity metrics. This shift from topology-constrained graphs to geometry-centered representations enables structural reasoning that is modality-agnostic and scalable.

The latent spaces learned by GTA-5 exhibit several notable properties. First, functional organization emerges naturally: pockets cluster according to Pfam families while retaining biologically meaningful heterogeneity. This partial, rather than absolute, segregation is critical. Perfect taxonomic clustering would merely reflect evolutionary similarity; instead, the observed neighborhood structure suggests that the model captures geometric and chemical compatibility that transcends strict family boundaries. Such behavior is particularly relevant for drug repurposing, where transferable structural features often occur across distantly related proteins. Second, analytically computed pocket properties—such as volume, aromaticity, and exposure—are recapitulated within the latent geometry despite not being used as direct supervision targets. Their alignment with the minimum spanning tree structure indicates that GTA-5 implicitly encodes physically meaningful features directly from raw three-dimensional data. This emergent interpretability bridges learned embeddings and classical pocket descriptors, suggesting that the model does not replace established geometric reasoning but internalizes it.

In the ligand space, structurally distinct scaffolds are positioned as neighbors when their geometric and compositional configurations are compatible. This behavior supports scaffold hopping within a purely embedding-based framework, without reliance on explicit fingerprint similarity. Importantly, this organization arises from spatial and semantic encoding alone, underscoring the importance of three-dimensional context in defining chemical proximity. A deliberate design decision underlying GTA-5 is the omission of explicit bond connectivity in the molecular representation. Conventional molecular graph neural networks enforce chemical topology as a primary structural prior. While appropriate for tasks centered on chemical reactivity or property prediction, this constraint limits representational flexibility when comparing fundamentally different structural objects such as ligands and binding pockets. By constructing proximity-based graphs dynamically from spatial neighborhoods, GTA-5 prioritizes geometric context over imposed connectivity. This choice introduces a controlled degree of structural “fuzziness,” allowing embeddings to reflect spatial compatibility rather than strict topological equivalence. The ligandome organization suggests that this abstraction does not erase chemical meaning but instead relaxes it sufficiently to enable cross-scaffold reasoning.

More broadly, GTA-5 establishes a framework for structural–chemical transferability. Embeddings derived from raw structural data can be interpreted as coordinates within a continuous compatibility manifold, where distances encode geometric and compositional similarity. Such manifolds are not merely descriptive but operational: they enable systematic navigation of pocketomes and ligandomes, quantitative neighborhood analysis, and hypothesis generation for repurposing or scaffold redesign. Unlike docking scores or energy functions, which evaluate specific ligand–target pairs, embedding distances provide a global structural context in which relationships can be explored at scale. The current study trains separate encoders for pockets and ligands to establish stable and interpretable latent geometries within each modality. However, the shared architectural formalism and bond-agnostic abstraction open a natural path toward unified cross-modality embedding spaces. Embedding ligands, pockets, and potentially peptides within a shared geometric language would enable bidirectional reasoning: pocket-to-ligand compatibility inference, ligand-to-pocket mapping, and structural interpolation across molecular classes. Such unification represents a critical step toward generative and transfer-based drug design strategies grounded in structural compatibility rather than target-specific heuristics.

Nevertheless, several limitations must be taken into consideration. The auto-encoding objective optimizes reconstruction fidelity of geometry and semantic labels but does not explicitly enforce task-specific objectives such as binding affinity prediction or synthesizability constraints. As with all unsupervised embeddings, downstream utility ultimately depends on the structure of the learned manifold and its alignment with functional outcomes. We intend to integrate in future work contrastive objectives and experimentally grounded validation loops to further calibrate embedding distances against biochemical measurements.

## 6 Conclusion

We present GTA-5, a geometry-centered graph transformer autoencoder that encodes protein binding pockets and small-molecule ligands with a unified three-dimensional point-cloud formalism. Deliberately representing structural objects through spatially contextualized chemical environments rather than assumed connectivity graphs given by atomic bonds, GTA-5 organizes large-scale pocket and ligand datasets into coherent and interpretable latent landscapes - shown here as a *pocketome* and *ligandome* constructed from the Protein Data Bank. Functional relationships emerge without explicit supervision, analytically defined pocket properties are implicitly recovered, and structurally distinct chemotypes converge when geometrically and semantically compatible. These results demonstrate that structural compatibility can be captured directly from raw three-dimensional data within a modality-agnostic embedding framework, allowing GTA-5 to support drug discovery applications including scaffold hopping in ligand-based virtual screening, QSAR/QSPR modeling using embedding-derived descriptors, and drug repurposing guided by pocket similarity. Beyond pocket and ligand encoding, this work establishes a representational foundation for structural–chemical transferability. Embedding diverse molecular entities within a shared geometric language lays the groundwork for unified latent spaces that can systematically relate pockets, ligands, and other molecular objects, facilitating cross-target reasoning and multi-modal molecular design strategies.

## Acknowledgements

This work benefited from a government grant (PPR Antibioresistance) managed by the Agence Nationale de la Recherche–Programme d’Investissements d’Avenir (ANR-20-PAMR-0011). BCC acknowledges funding support provided by the Pasteur-Roux-Cantarini fellowship of Institut Pasteur.

## Footnotes

1 https://www.ebi.ac.uk/pdbe/

2 http://pfam.xfam.org/

